# Predicting microsatellite instability from whole slide images using texture features

**DOI:** 10.1101/2025.03.30.646246

**Authors:** Nilus Swanson, Mauro A. A. Castro, A. Gordon Robertson, Ilya Shmulevich, Bahar Tercan

## Abstract

Identifying MSI in whole slide images (WSIs), one of the most widely used diagnostic imaging formats, is of great importance and in demand. In this study we employed color-based texture features to predict MSI on both a tile and sample based level. We found that within cohorts of hematoxylin and eosin (H&E) stained WSIs, texture morphology is able to predict MSI on a tile level with an AUC of up to 0.95 and on a sample level with an AUC of up to 0.98. This runs in contrast to other methods for predicting MSI in H&E WSIs which either utilized artificial intelligence based models, or achieved lower accuracy scores. Our results demonstrate that texture morphology is a significantly notable factor when it comes to identifying MSI in H&E WSIs, and should be used when constructing future models for MSI identification in a clinical setting.

## Introduction

Microsatellite instability (MSI) is a pattern of hypermutation that occurs at genomic microsatellites ^22^ and is caused by defects in the mismatch repair system (MMR) ^16^. Mismatch repair deficiency that leads to MSI has been well described in several types of human cancers, most frequently in adenocarcinomas from colorectal, endometrial, and gastric cancers. MSI is known to be a prognostic marker in gastric tumors which are able to be surgically removed ^16^; however, current clinical guidelines recommend MSI testing only for colorectal and endometrial cancers ^15^.

DNA contains repeated sequences of one to six nucleotides, which are known as microsatellites (MS). During DNA replication, MS sometimes contains errors in which these sequences have trouble properly repeating. Such errors are typically detected and repaired by a DNA repair system known as the mismatch repair (MMR). In the case of microsatellite instability (MSI), MMR proteins are missing, resulting in defects in repair, and subsequently increasing the chances of gene mutations and tumor development ^22^.

Several methods are available for detecting MSI. Next generation sequencing (NGS) uses targeted gene sequencing to identify correlation between MSI, MMR genes and the total number of mutations in the DNA, tumor mutational burden (TMB), in order to test for MSI presence in tumors. In 2017, NGS was approved for detecting MSI through Memorial-Sloan Kettering’s (MSK’s) IMPACT products ^1^. In 2018, NGS was approved for detecting MSI through Foundation Medicine’s (FMI’s) F1CDX.

Within normal and tumor tissue of patients, polymerase chain reaction (PCR) and capillary electrophoresis (CE) are the most common methods of MSI detection, using two single nucleotide repeat locations and three multi-nucleotide locations as parameters for the methods. Based on the number of sites demonstrating instability, the status of the tissue is determined to be MSS, MSI-L, or MSI-H. In this study, MSS and MSI-L were considered as one class, giving two classes of MSS and MSI (MSI-H) as is frequently done in clinical settings ^22^.

As missing MMR proteins are a factor in MSI, immunohistochemistry (IHC) is a method for detection of deficient mismatch repair (dMMR) and proficient mismatch repair (pMMR), which is when, respectively, one MMR protein is missing and when all four MMR proteins are missing. However, IHC results do not always correlate with the MSI status of the tissue being analyzed ^22^.

Single-molecule molecular inversion probes (smMIPs) are used in a method by the Academy of Sciences to detect MSI in a pan-cancer study ^7^, however, it only was able to accurately do so for colorectal, prostate, and endometrial cancers.

Finally, the other largely used method for MSI detection is through calculation by using a MANTIS score ^4^. MANTIS ^2,3^ is used for pan-cancer MSI detection in conjunction with ResNet-18, an 18-layer deep convolutional neural network (CNN), on histological images, and to the knowledge of us, is the only method to use histological images.

When testing for MSI, a common reason is lynch syndrome or hereditary nonpolyposis colorectal cancer, which is a disorder that increases the chances of contracting colon cancer. In terms of detection, lynch syndrome has been seen to be linked to dMMR tumors, and thus linked to MSI-H tissue. Lynch syndrome is difficult to detect when only using either a screening for MSI or IHC, so, instead, both must be used to detect it ^26^. As such, MSI detection is of great importance for detecting lynch syndrome.

Colorectal cancer (CRC) has been seen to be the cancer type most influenced by the presence of MSI; however, MSI has been seen to have effects on the prognosis of several cancer types, including gastric (GC), breast, prostate, cholangiocarcinoma, leukemia, bladder, ovarian, endometrial carcinoma (EC), pancreatic ductal adenocarcinoma (PDAC), follicular thyroid cancer (FTC), and adrenocortical cancers (ACC). Each needs to be analyzed and treated in various ways, partially based on their MSI status ^22^. CRC and GC are of particular note and typically fail to respond to immunotherapy in the way that other cancers do, unless there is MSI present in the tumor tissue ^31^.

In 2019, Kather *et al* ^31^ proposed a method for predicting MSI in H&E slides using deep learning. They trained several ResNet-18 models on respective datasets of The Cancer Genome Atlas (TCGA) colorectal FFPE and frozen, TCGA gastric FFPE, and TCGA endometrial FFPE. Each model was tested on a holdout validation set of their respective dataset. Additionally, Kather *et al*. tested their models trained on TCGA colorectal FFPE, TCGA colorectal frozen, and TCGA gastric FFPE against the Chancen der Verhütung durch Screening (DACHS) colorectal cohort ^13^. Kather *et al*. also tested their model trained on TCGA gastric cancer FFPE images against the Kanagawa Cancer Center Hospital (KCCH) gastric FFPE cohort ^6^. In 2021, Lee *et al*. ^15^ proposed a method for predicting MSI in H&E slides using an Inception-V3 neural network in colorectal, gastric, and endometrial cancers^15^. TCGA cohorts for colorectal, gastric, and endometrial cancers were used to train the respective versions of the model. In 2023, Lee *et al*. ^24^ proposed a Tensorflow deep learning method for prediction of MSI in gastric cancer (GC) ^24^. Two models were trained, one on TCGA GC FFPE tissue slides and the other on TCGA GC frozen slides. Both models were then tested on holdout sets of their respective cohorts. Additionally, the model trained on TCGA GC FFPE WSI was tested on an external validation FFPE cohort, made up of Asian patients from Seoul’s St. Mary’s Hospital (SSMH). In May 2024, Wang *et al*. ^17^ proposed a deep learning method for prediction of MSI in endometrial cancer. Wang’s method was both trained and tested on FFPE endometrial H&E stained WSIs from the TCGA cohort. When testing on TCGA endometrial, they separated the cohort into endometrioid carcinoma Type 1 (G1, G2) and endometrioid carcinoma Type 2 (G3). The accuracy, specificity, and/or balanced accuracy measures for all of the methods can be seen in Table 2.

**Table 1:** Most predictive features of each cohort.

| TCGA-CRC | TCGA-STAD | TCGA-UCEC | CPTAC-COAD | CPTAC-UCEC |
| --- | --- | --- | --- | --- |
| Correlation | Correlation | Correlation | Energy | Autocorrelation |
| Homogeneity | Homogeneity | Autocorrelation | Correlation | Correlation |
| Max Probability | Autocorrelation | Cluster Prominence | Max Probability | Sum of Squares |

**Table 2:** Accuracy Metrics for predicting MSI per-tile.

| Cohort | Cancer Types | AUC | Balanced Accuracy | Specificity | Precision (PPV) | Recall (NPV) |
| --- | --- | --- | --- | --- | --- | --- |
| TCGA-CRC | Colorectal | 0.84 (0.78 - 0.98) | 0.91 | 0.95 | 0.97 | 0.80 |
| TCGA-STAD | Gastric | 0.88 (0.83 - 0.98) | 0.93 | 0.93 | 0.93 | 0.92 |
| TCGA-UCEC | Endometrial | 0.95 (0.91 - 0.98) | 0.85 | 0.84 | 0.88 | 0.83 |
| CPTAC-COAD | Colon | 0.92 (0.88 - 0.95) | 0.82 | 0.80 | 0.80 | 0.84 |
| CPTAC-UCEC | Endometrial | 0.95 (0.94 - 0.97) | 0.90 | 0.88 | 0.87 | 0.90 |

In this study, we demonstrate a novel approach to predicting MSI status from H&E-stained whole slide images (WSI) for image tiles and for patients. Our method utilized grey level co-occurrence matrices to calculate Haralick features across red, green, and blue (RGB), hue, saturation, and value (HSV), and L-star, a-star, b-star (LAB) color spaces. By utilizing these highly interpretable features, this study was able to predict MSI from H&E-stained WSIs using cross validated RandomForest and XGBoost machine learning (ML) models. While many state of art methods utilize “black box” artificial intelligence (AI) based models for MSI detection, this study’s method utilizes specified mathematical formulation in order for detection of MSI in colorectal, gastric, and endometrial cancers.

## Methods

We obtained the formalin-fixed paraffin-embedded (FFPE) whole slide images (WSI) from the Kather *et al*. dataset ^31^. The images were of H&E-stained slides from (TCGA) project, for two different cancer types: colorectal adenocarcinoma ^30^ and stomach adenocarcinoma ^30^. Kather *et al*. ^31^ detected tumor regions, divided the WSI into 224 × 224-pixel tiles (i.e. tiles that are 112μm x 112μm), magnified to a resolution of 0.5 μm/px, normalized the color of the tiles using the Macenko method ^25^, and assigned a label of either MSI or MSS to each tile as previously described in ^5^. Figure 1 shows the flowchart of our approach.

**Figure 1.**
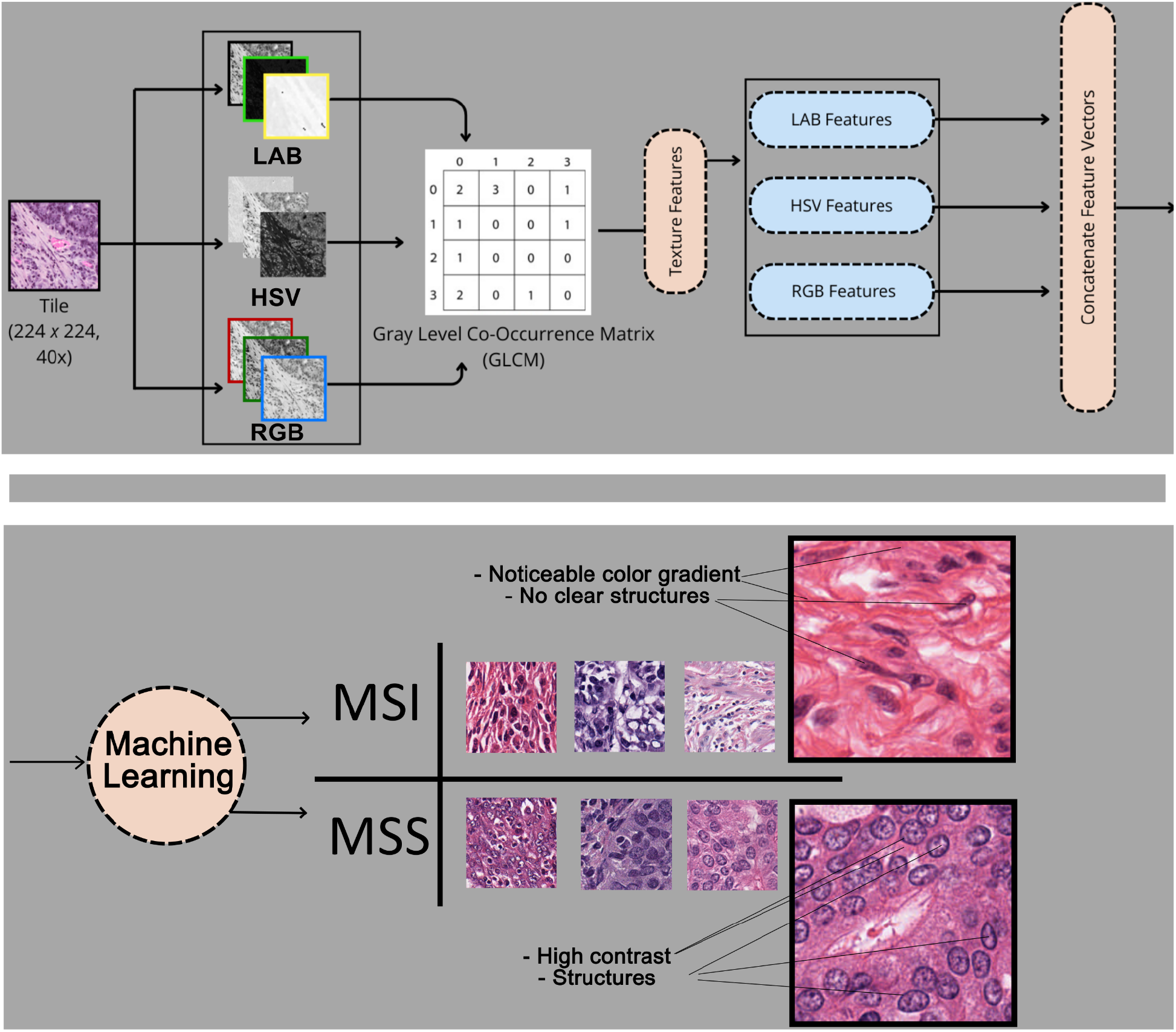
Flowchart of our approach. **Top panel**: Segmented tiles ^31^ are read in three different color spaces of RGB, HSV, and LAB. For each channel from those color spaces, four GLCMs are created, from which texture features are extracted. Those texture features are then concatenated into a single vector to represent a single tile. **Bottom panel**: The vector is passed onto a machine learning algorithm which then predicts if the tile is MSI or MSS ^31^.

Each tile was represented as a 3D matrix, with the first two dimensions representing the 2D tile image, and the third dimension representing the color channels of red, green, blue (RGB), hue, saturation, value (HSV), or L-star, a-star, b-star (LAB). For each of these nine 2D matrices, a separate matrix called a grey level co-occurrence matrix (GLCM) was created. This GLCM, *P*, was defined for each 2D image *I* as follows:

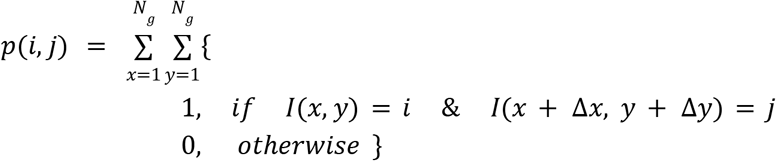

where (*i, j*) was the intensity value of each pixel at the location *I*(*x, y*). *N*_*g*_ represented the maximum intensity value of the color channel. (Δ*x*, Δ *y*) represented the angle offset at which co-occurrence was measured, which are equivalent to *θ*. The following angle offsets were used:

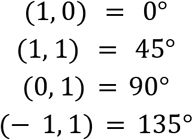

This means that for a single color channel of an image, four separate GLCMs were generated. For feature extraction, the tile images in this study were split into nine separate color channels, with each one generating 4 GLCMs, this would result in a total of 36 GLCMs per tile image, from each of which we extracted a number of features ^32^. The features are shown below.

*p* = grey level co-occurrence matrix (GLCM)

μ = mean of *p*

μ_*x*_ = mean of rows of *p*

μ_*y*_ = mean of columns of *p*

σ _*x*_= standard deviation of rows of *p*

σ *y*= standard deviation of columns of *p*

*N*_*g*_ = number of levels being measured by *p*

Marginal probability distribution 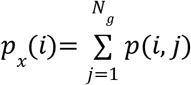

Marginal probability distribution 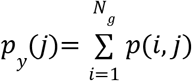

Marginal probability distribution 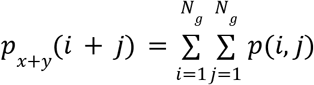

Marginal probability distribution 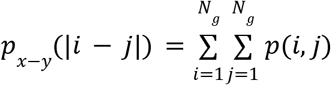

Contrast 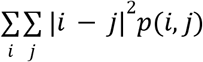

Correlation 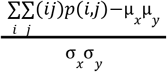

Dissimilarity 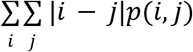

Energy 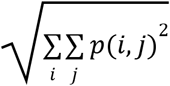

Homogeneity 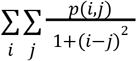

Angular Second Momentum (ASM) 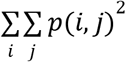

Autocorrelation 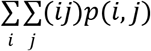

Cluster Prominence 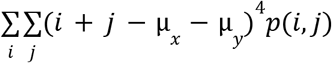

Cluster Shade 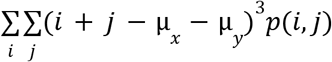

Entropy 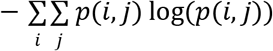

Maximum Probability 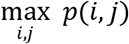

Sum of Squares 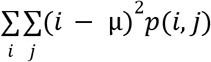

Sum Average 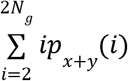

Sum Variance 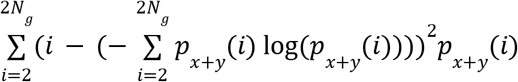

Sum Entropy 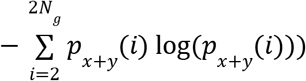

Difference Variance 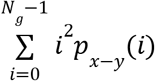

DIfference Entropy 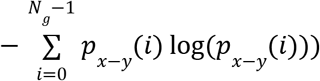

Normalized Inverse Difference 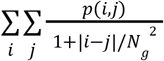

Normalized Inverse Difference Moment 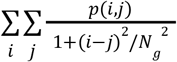

Trace 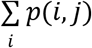

After we extracted these features, we utilized Random Forest in Python’s Scikit-Learn 1.5.2, as well as XGBoost 2.1.4, with cross validation and parameter tuning options to predict MSI. Each tile had a corresponding MSI/MSS label, as well as each sample it had come from. Therefore, once the per-tile predictions were done, we compared the ratio of MSI vs MSS tiles to the MSIMantis score, as labelled by ^29,30^ per sample to determine if a corresponding sample was MSI or MSS.

### Rationale

In our pipeline, each tile as segmented by ^31^ was separated into color spaces. From each color space, multiple GLCMs were created for each color channel, texture features were extracted and concatenated into a single vector per tile. Tiles were then run through machine learning and labelled as either MSI or MSS, as shown in Figure1.

From each cohort, we selected 18000 random tiles. We used 8000 tiles for training; and divided the remaining 10000 tiles into five test sets of n=2000 each. The train and test cohorts consisted of equal numbers of MSS and MSI samples whose labels are provided by ^16,31^. GLCMs were created as described in ^27^ for the color channels of red, green, blue (RGB), hue, saturation, value (HSV), or L-star, a-star, b-star (LAB) of each tile. 720 texture features were extracted from the GLCMs of each tile. Random forest and XGBoost with parameter tuning were used to create models for individual training cohorts for TCGA-CRC (colorectal), TCGA-STAD (gastric), and TCGA-UCEC (endometrial) FFPE WSIs, as well as CPTAC-COAD (colon) and CPTAC-UCEC (endometrial) frozen WSIs. Each model was then tested on randomly chosen stratified subsets of their respective cohorts, the results of which are shown in Table 4.

## Results

When we trained our models on the TCGA-CRC, TCGA-STAD, TCGA-UCEC, CPTAC-COAD, and CPTAC-UCEC cohorts, the per-tile reported values were area under curve (AUC) with a confidence interval (CI), balanced accuracy, specificity, precision, and recall, as shown in Table 2, with the corresponding Receiver Operating Characteristic (ROC) curves shown in Figure2. For TCGA-CRC, TCGA-STAD, and TCGA-UCEC the per-sample AUC and balanced accuracy are shown in Table 3, with the corresponding ROC curves shown in Figure3. For all five cohorts, the predictive features with the greatest weights are shown in Table 1.

**Table 3:** Accuracy metrics for predicting MSI per-sample.

| Cohort | AUC | Balanced Accuracy |
| --- | --- | --- |
| TCGA-CRC | 0.98 | 0.94 |
| TCGA-STAD | 0.98 | 0.97 |
| TCGA-UCEC | 0.95 | 0.93 |

**Table 4.** Accuracies of methods from literature.

| Study | Training Cohort | Testing Cohort | AUC or Specificity or Balanced Accuracy |
| --- | --- | --- | --- |
| Kather *et al.* 2019<sup>31</sup> | TCGA Colorectal FFPE | TCGA Colorectal FFPE | 0.77 AUC |
|  |  | DACHS Colorectal | 0.84 AUC |
|  |  | FFPE |  |
|  | TCGA Gastric FFPE | TCGA Gastric FFPE | 0.81 AUC |
|  |  | DACHS Colorectal FFPE | 0.60 AUC |
|  |  | KCCH Gastric FFPE | 0.69 AUC |
|  | TCGA Endometrial FFPE | TCGA Endometrial FFPE | 0.75 AUC |
| Schmauch *et al.* 2020 <sup>13</sup> | TCGA Colorectal FFPE | TCGA Colorectal FFPE | 0.82 AUC |
|  | TCGA Gastric FFPE | TCGA Gastric FFPE | 0.76 AUC |
| Valieris *et al.* 2020 <sup>12</sup> | TCGA Gastric FFPE | TCGA Gastric FFPE | 0.81 AUC |
| Krause *et al.* 2021 <sup>10</sup> | TCGA Colorectal FFPE | TCGA Colorectal FFPE | 0.742 AUC |
| Lee *et al.* 2021 <sup>15</sup> | TCGA Colorectal FFPE | TCGA Colorectal FFPE | 0.861 AUC |
| Lee *et al.* 2023 <sup>24</sup> | TCGA Gastric FFPE | TCGA Gastric FFPE | 0.902 AUC<br>0.833 Specificity<br>0.862 Balanced Accuracy |
|  |  | SSMH FFPE | 0.874 AUC<br>0.806 Specificity<br>0.802 Balanced Accuracy |
|  | SSMH and TCGA Gastric FFPE | TCGA Gastric FFPE | 0.756 Specificity<br>0.847 Balanced Accuracy |
|  |  | SSMH FFPE | 0.887 Specificity<br>0.895 Balanced Accuracy |
| Wang *et al.* 2024 <sup>17</sup> | TCGA Endometrial G1,G2 FFPE | TCGA Endometrial G1,G2 FFPE | 0.94 Balanced Accuracy |
|  | TCGA Endometrial G3 FFPE | TCGA Endometrial G3 FFPE | 0.84 Balanced Accuracy |

**Figure 2:**
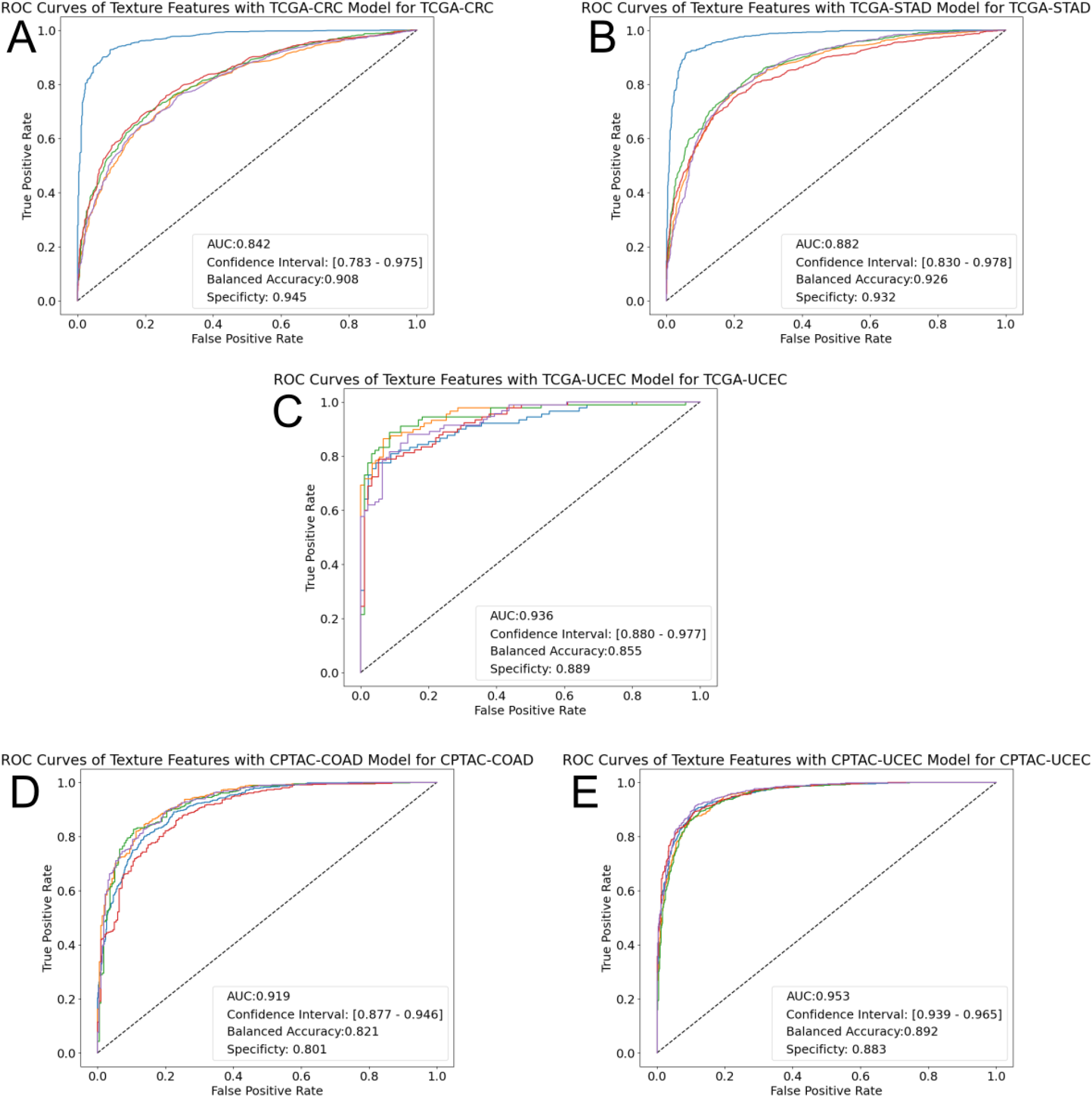
ROC curves of MSI prediction across each cohort. Each curve shows one test split. The reported AUC is the mean average across the splits. **A**: TCGA-CRC (colorectal) **B**: TCGA-STAD (gastric) **C**: TCGA-UCEC (endometrial) **D**: CPTAC-COAD (colon) **E**: CPTAC-UCEC (gastric)

**Figure 3:**
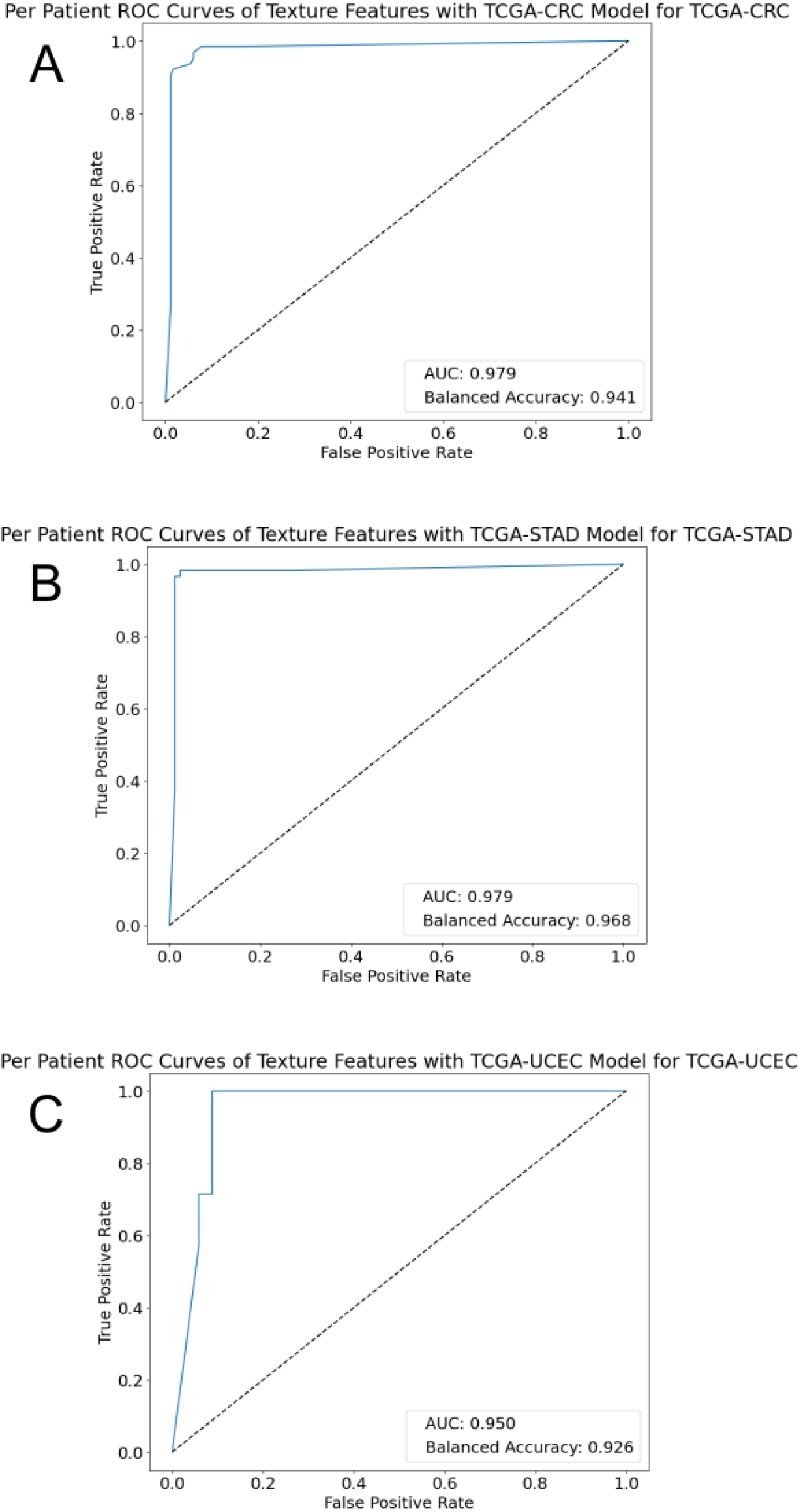
ROC curves of per-patient MSI prediction model. **A**: TCGA-CRC (colorectal) **B**: TCGA-STAD (gastric) **C**: TCGA-UCEC (endometrial).

## Discussion

In this study, we utilized texture properties of whole-slide H&E stained images in order to predict microsatellite instability of colorectal, gastrointestinal, and endometrial cancers using cross validated random forest and XGBoost models. Our results suggest that the texture morphology of WSIs can be used to predict MSI within all three cancer types (colorectal, endometrial and gastric) when trained on separate WSIs of the same cancer type. All of the TCGA WSIs trained and tested on were FFPE WSIs. All of the CPTAC WSIs trained and tested on were frozen WSIs.

While this is not the first study to indicate a distinction in MSI and MSS detectable through H&E WSIs, to our knowledge, it is the first to do so using solely texture morphological features. Other studies on detecting MSI within H&E samples primarily utilized artificial intelligence (AI) based methods. The results of these studies, and the comparable metrics to our study, are shown in Table 2. While the other studies reported an AUC value for most of their prediction results, Lee *et al*. ^24^ did not do so for two of their test cohorts, and Wang *et al*. ^17^ did not report an AUC for any of their test cohorts. As such, the balanced accuracy was used as the chosen metric of comparison between these studies and our study.

Our study, when trained and tested on TCGA colorectal cancer WSI’s, reported an AUC value of 0.84 (0.78-0.98), exceeding that reported by Kather *et al*. ^31^, Schamuch *et al*. ^13^, and Krause *et al*. ^10^ when tested on the same cohort.

Our study, when trained and tested on the TCGA gastric cancer cohorts an AUC of 0.88 (0.83-0.98), exceeding the results reported by Kather ^31^, Schamuch ^13^, and Valieries ^12^, when tested on the same cohort, but falling just below those reported by Lee ^24^. However, when the balanced accuracy and specificity of this study for predicting MSI in TCGA gastric cancer cohort was 0.93 and 0.93 respectively. In contrast, Valieries reported balanced accuracy and specificity scores of 0.862 and 0.833. This suggests that texture-based features are strong predictors for correct identification of MSI and MSS, as also evident by the positive and negative predictive values as shown in Table 2, reported as 0.93 and 0.92 respectively for TCGA gastric cancer.

Kather *et al*. ^31^ and Wang *et al*. ^17^ both trained at least one model on TCGA endometrial FFPE WSIs. Kather *et al*. predicted TCGA endometrial cancer MSI with an AUC of 0.75. Wang *et al*. predicted TCGA endometrial in two separate cohorts, being separated in TCGA endometrial G1 and G2 and TCGA endometrial G3. These differentiate by percentages of non-squamous areas, with G1 having “no more than 5% solid non-squamous areas,” G2 having 6-50% non-squamous areas, and G3 having “more than 50% solid non-squamous areas” ^19^. Squamous areas of the Amant *et al*. ^19^ found that tumors “upgraded from grade 1 to 2, or from grade 2 to 3, if there is striking cytological atypia”, which suggests that there could be difference in texture morphology between endometrial cancer grades. Wang *et al*. reported a balanced accuracy of 0.94, when trained and tested on TCGA endometrial G1&G2 and 0.84 when trained and tested on G3. Our study, when trained on the TCGA endometrial cohort and tested on a holdout test set of TCGA endometrial cancer, reported an AUC value of 0.95 (0.91-0.98), with a corresponding balanced accuracy and 0.85. While our study did not account for the differences in endometrial cancer grades, as Wang *et al*. did, and reported a noticeably high AUC, the balanced accuracy fell below that reported by Wang *et al*.

Notably, both of the CPTAC-COAD and CPTAC-UCEC cohorts are not FFPE WSIs, but instead frozen WSIs. As such, they are subject to warping during the freezing process, unlike when a sample is enclosed in paraffin wax. Regardless of that, it would seem that the models trained on frozen WSIs were able to achieve greater AUC scores than those trained on FFPE WSIs, with the exception of TCGA-UCEC.

In the current state of histological, and biological research in general, it is very tempting to utilize AI-based models to perform analyses on multitudes of samples. After all, a highly trained and finely tuned model analyzing an image could detect patterns that the human eye has yet to. This study highlights n that texture is a highly effective set of features to implement into future models for higher accuracy MSI prediction, as well as possibly many other morphological features of H&E WSIs, as evident in Figure 1. As can be seen in those example tile images, texture features, such as a distinct change in contrast, are visible to the naked eye. Such features being noticeable further lends itself to the recorded observations of this study, that texture morphology is a notable feature in distinguishing MSI from MSS within histopathological whole slide images. As such, this study suggests that future models for usage within a clinical setting should make use of texture features for identifying MSI.

## Author Contributions

I.S. conceptualized the study, NS implemented the analyses and visualizations and wrote the initial manuscript draft. B.T., I.S., M.C., G.R. provided guidance, I.S. and B.T. administered the project, B.T., N.S., M.C., G.R. did manuscript reviewing and editing.

## Data and Code Availability

The TCGA data used in this study is openly available in the GDC data portal (https://portal.gdc.cancer.gov). The TCGA-CRC and TCGA-STAD tile segmentations used in this study are openly available at https://zenodo.org/records/2530835. The code for the method used in this study is openly available on Github (https://github.com/IlyaLab/TextureAnalysis).

## Acknowledgments

We would like to dedicate this paper to the memory of Ilya Shmulevich, who initiated this study and co-mentored Nilus Swanson with Bahar Tercan. Nilus Swanson and Bahar Tercan are grateful to the Institute for Systems Biology.

